# CRISPR-GNL: an improved model for predicting CRISPR activity by machine learning and featurization

**DOI:** 10.1101/605790

**Authors:** Jun Wang, Xi Xiang, Lixin Cheng, Xiuqing Zhang, Yonglun Luo

## Abstract

**Motivation:** The CRISPR/Cas9 system has been broadly used in genetic engineering. However, risks of potential off-targets and the variability of on-target activity among different targets are two limiting factors. Several bioinformatic tools have been developed for CRISPR on-target activity and off-target prediction. However, the general application of the current prediction models is hampered by the great variation among different algorithms.

**Results:** In this study, we thoroughly re-analyzed 13 published datasets with eight regression models. We proved that the current model gave very low cross-dataset and cross-species prediction outcome. To overcome these limitations, we have developed an improved model (a generalization score, GNL) based on normalized gene editing activity from 8,101 gRNAs and 2,488 features using Bayesian Ridge Regression model. Our results demonstrated that the GNL model is a better general algorithm for CRISPR on-target activity prediction

**Availability and implementation:** The prediction scorer is available on GitHub (https://github.com/TerminatorJ/GNL_Scorer).

**Contact:** J.W. or Y.L.

**Supplementary Information:** Supplementary data are available at Bioinformatics online.

## INTRODUCTION

CRISPR/Cas9 is the adaptive immunity system in bacteria and most archaea (Horvath and Barrangou, 2010; Koonin and Makarova, 2009), which was first harnessed for programmable and precision gene editing in 2012. The CRISPR/Cas9 gene editing system is comprised of two key components: a small guide RNA (gRNA) and a Cas9 endonucleases (Deltcheva, et al., 2011; Martin Jinek, et al., 2012). The gRNA is a chimeric RNA molecule of tracrRNA and crRNA (Ran, et al., 2013) that guides the Cas9 protein to the target site in the genome. Another important feature of the CRISPR-Cas9 gene editing system is its dependence on a high conserved DNA motif, also known as the protospacer adjacent motif (PAM). The PAM-dependent targeting of CRISPR is able to distinguish self and non-self-DNAs. During CRISPR/Cas9-mediated gene editing, a targeted double-stranded DNA break (DSB) is firstly introduced. DSBs are subsequently repaired by the DNA repair pathways in mammalian cells: nonhomologous-mediated end joining (NHEJ) and homology-directed repair (HDR). Thus, the performance of CRISPR gene editing is decided by four layers, (1) selection of the best CRISPR gRNA target site (or guide sequences); (2) an efficient delivery of the CRISPR components to cells/host; (3) introducing a precise DSB; and (4) DSB repaired by NHEJ or HDR.

Selecting a target site with high on-target activity and low off-target is substantially important for gene editing. Previously, we have discovered that the gene editing activity of CRISPR-Cas9 in mammalian cells is affected by several factors, such as the secondary structure and chromatin accessibility of the guide sequences (Jensen, et al., 2017). To select gRNAs with high activity, we also developed a dual-fluorescence system for experimentally evaluating gRNA activity in cells (Zhou, et al., 2016). Previous results from us and other research groups consistently reveal that the CRISPR gRNA activities are highly variable. Thus, several *in silico* gRNA design web tools and algorithms have been developed to facilitate CRISPR design and applications.

Three types of CRISPR design software have been developed based on both experimental and simulated data: (*i*) alignment-based, where the CRISPR guide sequences were retreated based on mapping PAM sequences in the genome; (*ii*) hypothesis-driven, in which gRNA activity score is given based on the specific features such as GC content. (*iii*) machine learning based, where gRNA efficiency score is predicted by models trained with big datasets (Chuai, et al., 2017). An increasing number of CRISPR gene editing data are generated by different laboratories around the world. Big data-driven leaning based methods have become the keys in CRISPR gRNA activity prediction. Also, a growing number of features have been discovered to be of importance for gRNA activity. Thus, featurization of the gRNA sequences has been widely used to improve the prediction accuracy, such as SSC (Xu, et al., 2015), gRNA Designer (Doench, et al., 2016), gRNA Scorer (Chari, et al., 2015), CRISPR-DT (Zhu and Liang, 2018). DeepCas9 (Xue, et al., 2018), CNN_std (Lin and Wong, 2018) and CRISPRcpf1 (Kim, et al., 2018) utilized the convolution neural network (CNN) to predict the gRNA activity based on the automatic recognition of sequence features, resulting in high performance in both Cas9 and cpf1 protein.

However, limitations exist when applying these models for gRNA activity prediction, such as the high variability among different species and datasets. Species-specific software were developed, such as fryCRISPR for *Drosophila* (Gratz, et al., 2014), CRISPR-P for plant (Lei, et al., 2014), and CRISPRscan for zebrafish (Moreno-Mateos, et al., 2015). Machine learning based software, such as CRISPOR, have been applied in different species (Chuai, et al., 2017). However, the prediction models had a great bias and limitation for species generalization (Guo, et al., 2018). A benchmark comparision has been done in CRISPOR showing that the generalization of different scores are extremely low (Haeussler, et al., 2016). So, the heterogeneity of different species needs to be explored. In addition, batch effects and gRNA activity detection methods used different experiments have not been taken into consideration.

In this study, we applied 11 published datasets across different experiments, cells, and species, to compare the performance of each dataset using the same model and evaluation index. Eight models were used for data training to evaluate the data quality and the measurement impact of the experiments. We identified significant variations and difference by the Spearman correlation test. Our results show that the software trained by data from one species is not appropriate for the others. Additionally, an improved model, GNL Scorer (https://github.com/TerminatorJ/GNL_Scorer), was proposed based on data selection and feature collection and trained with two datasets (12340 gRNAs) (Hart, et al., 2015).

## MATERIALS AND METHODS

### gRNA efficiency data source

We initially retreated 14 datasets from different experiments and five species: human, mouse, zebrafish, *Drosophila*, *Ciona intestinalis* (**Table1**). The HEL dataset was used as the independent validation set together with Z_fish_GZ, Z_fish_VZ, Ciona, *Drosophila*, *C. elegans* which were not used to build any other model. We used human knock-out efficiency datasets from six experiments used nine cell types (HCT116 (Hart, et al., 2015), Hela (Hart, et al., 2015), HEK293T (Chari, et al., 2015), Hl60 (Wang, et al., 2014), NB4 (Doench, et al., 2014), TF1 (Doench, et al., 2014), MOLMB (Doench, et al., 2014), A375 (Doench, et al., 2016), HEL (Labuhn, et al., 2017)). As for the mouse dataset, we initially used MEsc cell dataset (Koike-Yusa, et al., 2014) and EL4 dataset (Doench, et al., 2016). Three types of zebrafish datasets were used: Gagnon et al. (Gagnon, et al., 2014), Moreno-Mateos et al. (Moreno-Mateos, et al., 2015), and Varshney et al. (Varshney, et al., 2015). We also used the Ciona (Gandhi, et al., 2016), *Drosophila* (Ren, et al., 2014), and *C. elegans* datasets (Farboud and Meyer, 2015) as testing datasets. The MEsc and Hl60 datasets were bimodal distributed, which were not included in BRR model training to minimize the impact on generalization.

For comparison of generalizations for cross-species prediction, the datasets were divided into four categories according to species. Human datasets were from all the human cell type data except HEL. Mouse datasets were separated from the Doench V1 datasets (Doench, et al., 2014), which was trained the gRNA Designer (Rule set I) by 959 gRNA, and MEsc dataset with 949 gRNA from Koike-Yusa et al. (Koike-Yusa, et al., 2014). *Ciona intestinalis* and *Drosophila* datasets were used as previously described.

### Normalization of datasets

For better comparability between different datasets, we normalized all the single datasets with batch-effect to the range from −4 to +4. And the dataset of *C. elegans* which could not be normalized to this standard was removed in the combining normalized single dataset. The tool we used to normalize the data was SPASS 20.0. All the 11 datasets (MEsc and Hl60 datasets excluded) were combined into a single dataset (all-in-one, AIO) for testing the influence of model performance in the same standard of features and algorithms.

All the 10 normalized datasets were used for training when applied to select the best dataset for generalized prediction. We randomly split each of them: 80% for training and 20% for testing. There are six datasets that are used as the validation datasets for comparing the performance of different algorithms. These six datasets were not used to build the machine learning model for other software. For evaluation of the generalized performance, we selected the human HEL, *Drosophila*, *C. elegans*, zebrafish, and Ciona as the independent validation datasets.

### Predictive models

All prediction models used in this study were based on regression. Eight regression models were used: (i) Gradient-boosted regression tree (GBRT), (ii) Decision tree (DT), (iii) Linear regression (Linreg), (iv) L2-regularized linear regression (L2reg), (v) L1-regularized linear regression (L1reg), (vi) Bayesian Ridge Regression (BRR), (vii) Random forest (RF), (viii) Neural network (NN). All these models were applied by using the scikit-learn package in python. We set the learning rate of 0.01 in GBRT model. The alternative max depth was set to [5, 10, 100, 1000] for grid search in the DT model. We provided alpha = [1, 3, 5, 7, 9, 10] to select grid in the L2reg model and alpha = [0.1, 0.5, 0.75, 1, 10] for grid search in L1reg model. As for the Lingre, BBR and RF models, default settings were used. For the NN model, we used two hidden layers with 100 neurons each. We use the 10-fold cross validation for selecting the hyper-parameter setting of each candidate model.

### Featurization

Three sets of features were evaluated in this study. Feature set I contains 627 features, and is partially derived from the Doench features (Doench, et al., 2016). The feature set I includes six parts of features. (*i*) 604 features of “one-hot” encoding of the nucleotide. There are two subsets in this category: position-dependent and position-independent. And each category applies to the one nucleotide and pairwise nucleotide. Such as “_nuc_pd_Order2” consisted of e.g. AA_1/AT_1/AG_1, and “_nuc_pd_Order1” consisted of e.g. A_1/T_1/G_1/C_1. (*ii*) 3 GC features, which consists of GC count, GC count < 10, GC count > 10. (iii) 16 features of the two nucleotides flanking the NGG PAM in the 5’ and 3’. (iv) Seven thermodynamic features. We calculated four thermodynamic features as Doench et al. (Doench, et al., 2016) using the “Tm_staluc function” in Biopython package. All these features above were derived from the 30mer of target sequences.

The feature set II is a combined set of 2488 features. In addition to those included in the feature set I, we added three extra type of features. (i) Four features of a consecutive of five nucleotides, such as AAAAA/GGGGG. (ii) 1856 features of three nucleotides with “position-dependent” and “position-independent”, such as ACG/AGG and ACG_1/ACT_2. (iii) One feature of delta G. The delta G of seed spacer sequence was calculated by hybrid-ss-min program in the OligoArrayAux package (Zuker, 2003). Note that, (i) and (ii) used the 30mer context sequence. And (iii) used the 20nt guide sequences.

Feature set III contains 2701 features. On top of feature set II, we added three additional feature sets: (i) 200 sequence conservation features, (ii) 10 sequence disorder status features as described by Chen et al. (Chen, et al., 2017), and (iii) Three epigenetic features: chromosome accessibility, CTCF and H3K4me3 from the ENCODE (Consortium, 2004). Position specific iteration BLAST (PSI-BLAST) was used to search the homologous protein against the query protein. We used the NR (non-redundant) database to retrieve the homologous proteins. And the PSSM matrix was calculated by SPOT-Disorder (Heffernan, et al., 2015). The 30mer context sequences are converted to the corresponding amino acid (AA) sequences (10 aa). If the resulting length of the amino acid sequences was shorter than 10, we inserted “X” for the lacking AAs. After generating the PSSM matrix, the 10-amino acid target protein was encoded as 200 (20×10) PSSM features. The disorder status was calculated by SPOT-Disorder (Heffernan, et al., 2015). Each 10-AA sequence was given a disorder probability in this residue. In total, 10 disorder features were included.

### Feature selection

The 2701 features were used to represent the factors that may influence CRISPR gene editing efficiency. However, not all of them would contribute equally. Thus, we used the mRMR method (Peng, et al., 2005) to rank and select features of importance. According to the relationship between the features and their target classes, each feature was ranked in a descending order and generated a feature list called MaxRel. The MaxRel list was generated as described below. Firstly, we calculated the mutual information (MI) to access the correlation between two variables, which are two features among the 2701 features. Using a probabilistic density function P(x), P(y) and P(x,y), P(x,y) was denoted as the joint probabilistic density of P(x) and P(y):

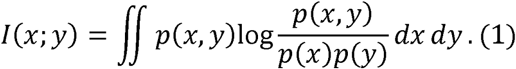

Next, the highest value of MI to the target class c was calculated. {x_i_,i=1,…,m} was the feature set of m features in S, required to have the largest value of MI to the target class. The MI was denoted as D here.

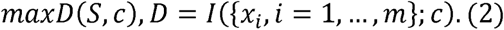

Third, the minimal redundancy between each feature pair was calculated. x_j_ was selected feature in the whole feature set S, and x_j_ is the feature need to be selected. When we calculate in the constrain of (2) and (3) is called “minimal-redundancy-maximal-relevance” (mRMR) (Ding and Peng, 2005).

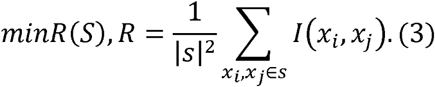

For each x_j_ in the reset of feature in the S, D-R is calculated, Then the x_j_ is transferred to x_i_. When all the x_j_ is turn to x_i_, the whole procedures will be stopped.

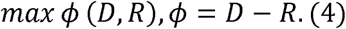

After completing the calculation as described above, we generate list as in (5). And we will also generate a serial of feature sets like S1, S2, S3 in the S. Then, each sub feature set was used to train the BBR model and measured by a corresponding evaluation value (Chen, et al., 2017). This method was called IFS (incremental feature selection).

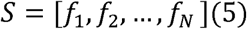

To select the best combination of features, we use the value of Spearman Correlation when training the model in the way of IFS. The best BBR was trained and got the best Spearman value of 0.49 when use 1839 features.

### Model evaluation

We used Spearman Correlation to assess the regression model, the Spearman Correlation could make use of the real-valued scores better than classification evaluation (Fusi, et al., 2015). The Classification-based algorithm discards the real-valued assay measurement, and converts them to binary active/inactive. Thus, it is unable to capture the more nuanced information available in the data. The Spearman Correlation conducted using the *Scipy* package used in Python.

## RESULTS

### Data normalization can alleviate batch-effect among different CRISPR activity datasets

We used 13 broadly used datasets (summarized in Table 1) with the objective of developing a new model with good performance and generalization. The Doench dataset (Doench, et al., 2016) was divided into 2 versions (V1 and V2) based on how CRISPR gRNA activity was evaluated (resistance assay or flow cytometer). Since all the 13 different datasets are from different studies and species, we first evaluated the data normality and found that not one was normally distributed (**Figure S1**). Thus a data normalization step was applied (see method) and after normalization, the *C. elegan* dataset (Farboud and Meyer, 2015) was removed due to failure in data normalization. We normalized the remaining 12 datasets to the gRNA activity range from −4 to 4 (**Figure S2**). Then we combine them as a new dataset called “normalized all-in-one (AIO) dataset”.

**Table 1:**
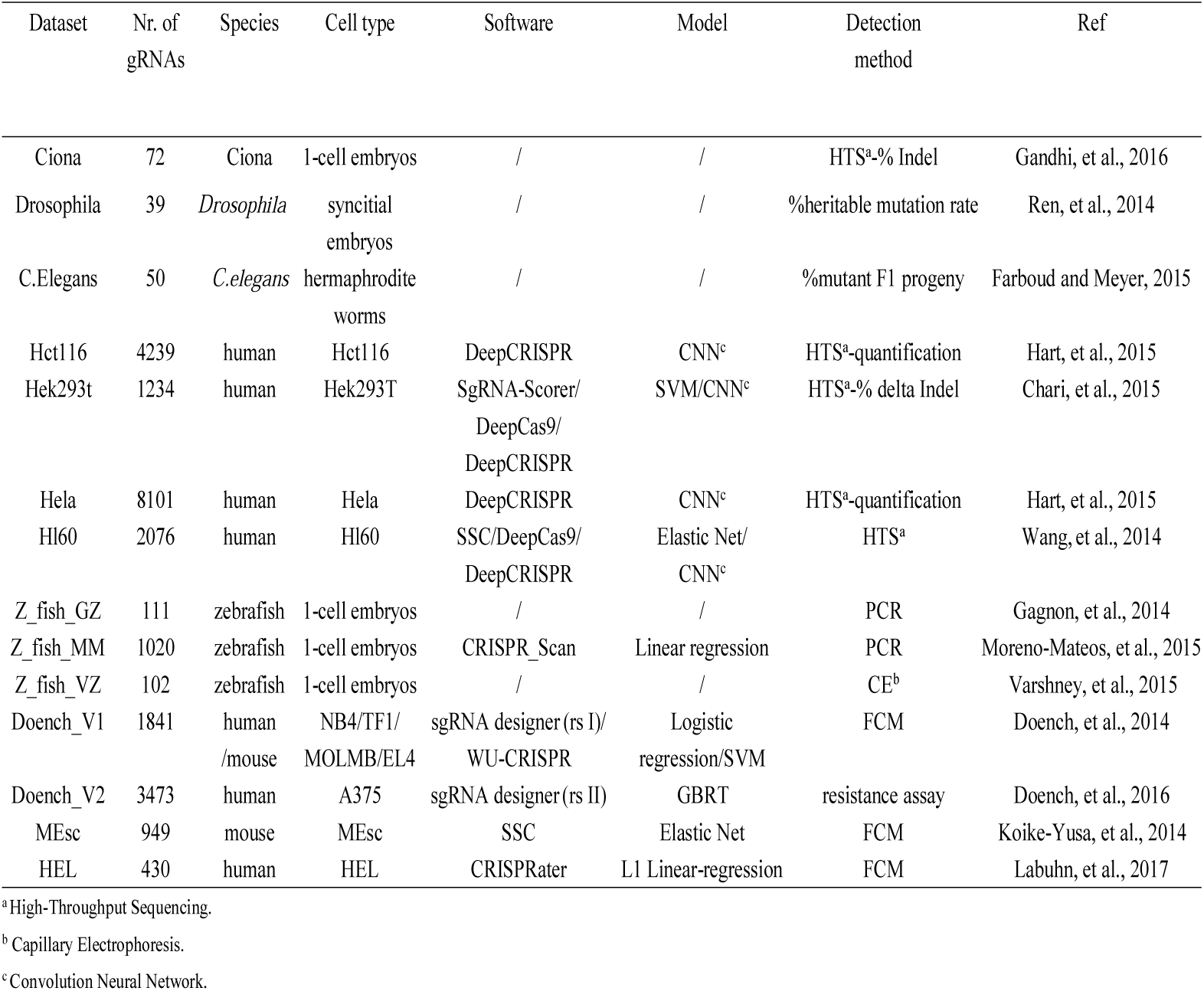
Summary of CRISPR activity datasets used in this study

To evaluate whether data normalization can alleviate batch effect, we train the normalized and non-normalized Doench (V1 + V2) and AIO datasets using the GBRT (Gradient Boosted Regression Tree) model, a state-of-art model used in gRNA Designer (rule set II) in the feature set I. Our results showed that the non-normalized AIO dataset could not improve the Spearman Correlation by just adding the data without considering the batch-effect of different datasets from different experiments (Figure 1). The Spearman Correlations for normalized AIO dataset were significantly improved (p value = 0.045) compared to the non-normalized AIO data, while no difference was found for the normalized and non-normalized the Doench datasets (Figure 1). However, even with the combined normalized AIO dataset, the GBRT Spearman Correlation value is significantly lower (p value = 0.044) compared to the Doench datasets. This results collectively suggested that there is a strong batch effect among the different datasets. Most importantly, it indicates that algorithms developed with a specific training dataset may have a strong bias when used to predict the other datasets.

**Figure 1:**
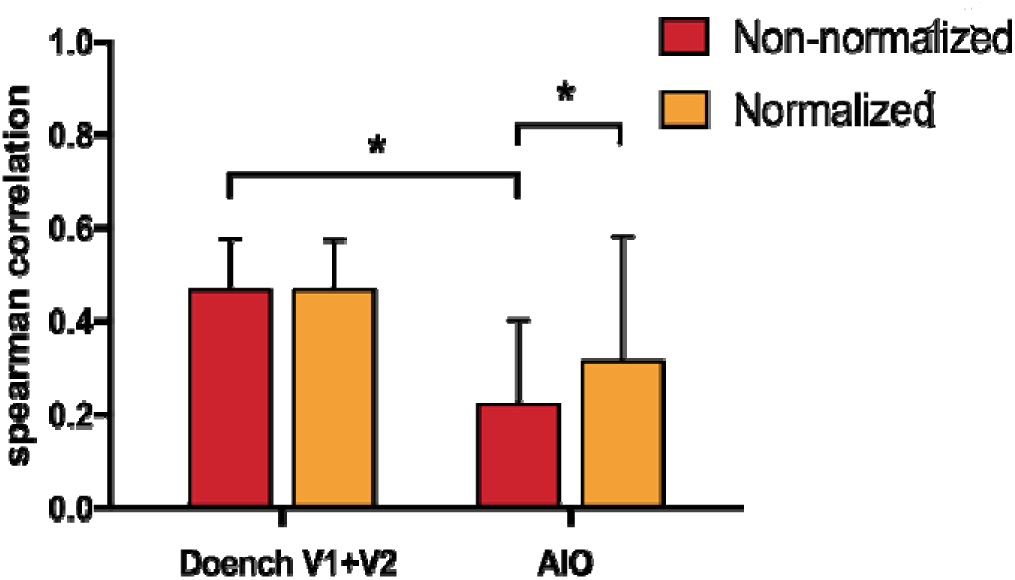
Doench dataset and AIO dataset trained by GBRT (Gradient Boosted Regression Tree) Error bars show the s.d. across sgRNA with a 10-fold approach. Doench dataset outperforms than AIO from left to right, Significance results are from paired t-test, *Asterisk*(*) indicates a *p* value less than 0.05.

### Selection of datasets with the best prediction outcome

To select good datasets that give the better general prediction of gRNA activity, we used 10 normalized datasets and randomly divided each dataset: 80% for training and 20% for testing. And we trained 8 models (see methods) with the 80% training dataset. The MEsc and Hl60 were not included due to bimodule distribution of the dataset (**Figure S1**). We next chose the model with the highest Spearman Correlation for each specific dataset (**Figure S3**), and evaluated the generalization of the remaining 20% testing datasets in all 10 groups. We found that Hela, HCT116 and Doench V1 datasets had the best performance in overall gRNA activity prediction (Figure 2a, b). The activity of the gRNAs in the Hela and HCT116 datasets were measured by a quantized sequencing detected method (Hart, et al., 2015). In addition, the gRNA activity datasets generated by sequencing give the best generalization performance compared to the rest of the datasets generated by other methods. This indicates that the dataset detected by sequencing-based method could have great generalization ability. Consistent with our finding, the Hela and HCT116 datasets had a “cleaner” supervisory signal for machine learning the same as FC to RES dataset (Fusi, et al., 2015).

**Figure 2.**
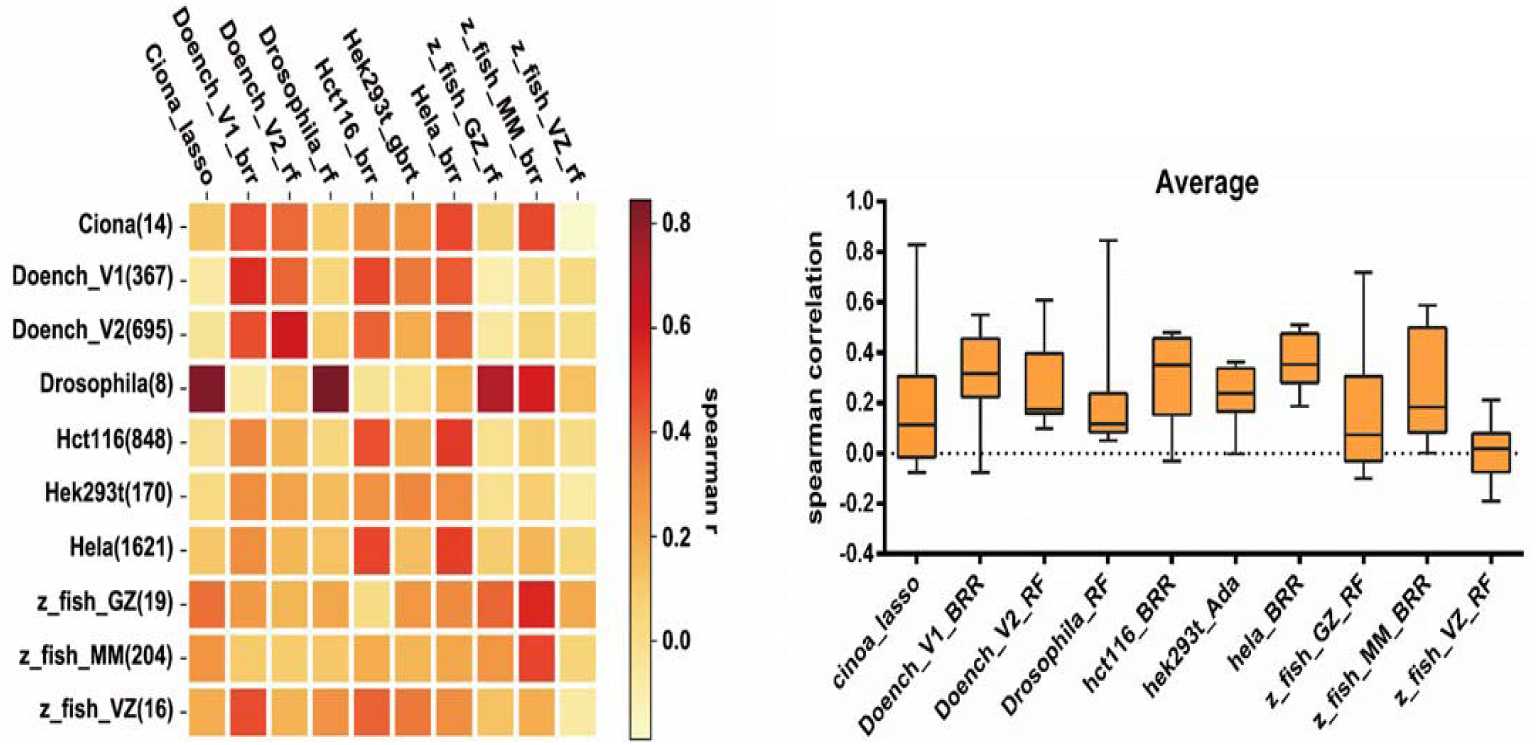
The performance of each best model in specific dataset. a) The x axis is the best model for specific 80% train dataset, the y axis was the 20% test datasets. The heatmap shows from the color of yellow to crimson. The darker color means the higher Spearman correlation. b) The mean value of Spearman correlation for each model trained by 80% of specific dataset and predicted by the remained 20% datasets.

Next, we investigated whether the combination of best-performing datasets could further improve model performance compared to the best performing Hela dataset, which we shown that has a great generalization gRNA activity prediction. All three datasets used the BRR as their best performance model (**Figure S3**). We next trained BRR model with four sets of data: (1) Hela dataset; (2) Hela+Doench_V1 datasets, (3) Hela + HCT116 datasets, (4) Hela + HCT116 + Doench_V1 datasets. The result show that the combined HCT116 and Hela dataset has a higher generalized prediction power than the other datasets combination (Figure 3). and similarly well as compared to the Hela dataset along.

**Figure 3.**
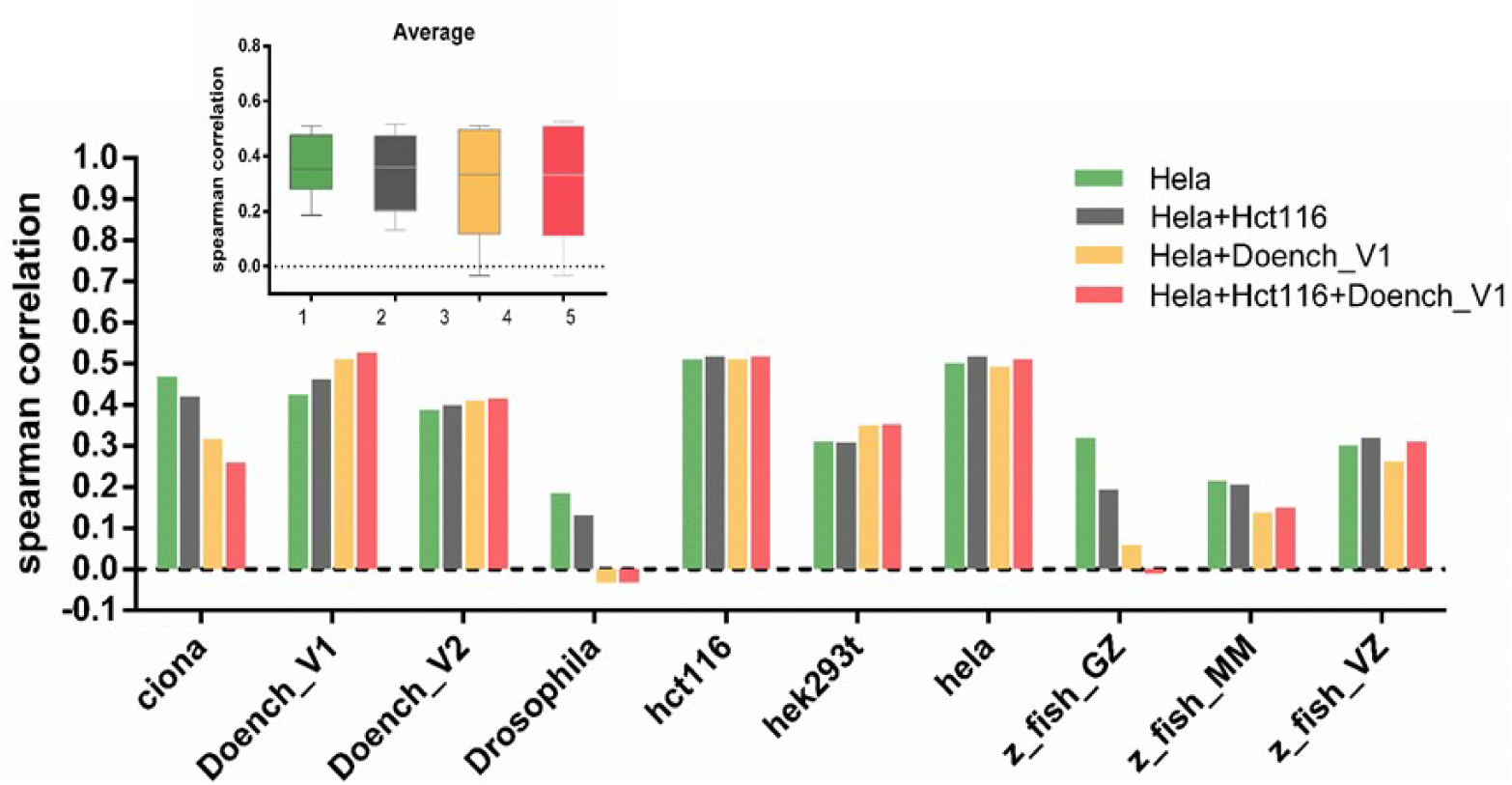
Combination of the normalized datasets with the highest Spearman correlation. All the datasets are trained by BRR (Bayesian Ridge Regression), which perform the best among each dataset as mentioned before. Here we just split the best three datasets that have the best generalization. Due to the other datasets were less likely to perform well than these. x-axis shows the 20% testing datasets for each one, the Spearman correlation value is the average score of 10-fold cross validation. The thumbnail shows the mean value across all these test datasets for four combined models

### Datasets from different species have pool generalization of gRNA activity prediction

As observed above, we next investigated whether models trained by one specific species could be used for gRNA activity prediction in other species. Currently, although all *in silico* CRISPR gRNA design web tools are defined for each species mainly due to their genome specificity. However, the prediction of gRNA activity is not defined as species-specific. To validate that, we grouped the normalized datasets into five species: human, mouse, zebrafish, *C. elegans*, *Drosophila*. Firstly, we trained each regression models using the combined data (the training dataset, 80%) and selected the best performing model for each combined dataset (**Figure S4**). Next, we evaluated the accuracy of gRNA activity prediction of species-specific datasets (the testing dataset, 20%s) using the best performing model. Our results showed that each best performing model trained with a species-specific dataset can well achieve self-prediction (Figure 4). However, all models have a pool generalization of cross-species gRNA activity prediction. It was noted that the models trained with human or mouse datasets are more effective in cross-species prediction.

**Figure 4:**
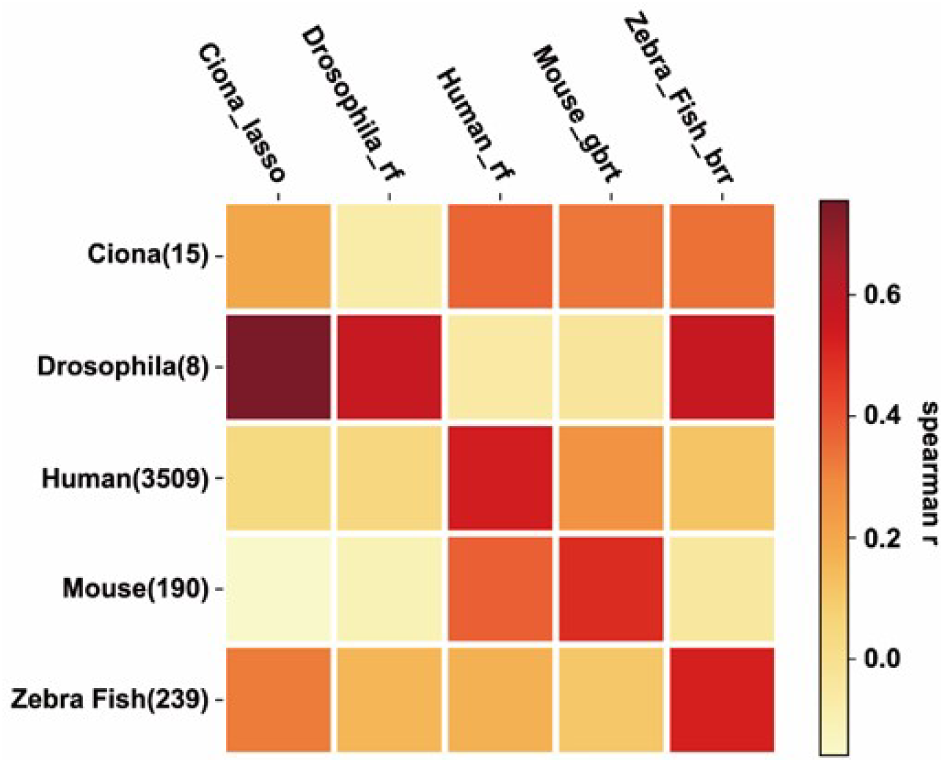
The generalization of the models trained by different species. The rows represent the best model of specific species trained by 80% of each dataset, the optimized model is tested by 20% dataset remained from each species dataset as shown in the columns.

### Evaluate features importance for gRNA activity prediction

Sequence specific, thermal dynamic, and epigenetic features affect CRISPR gRNA activity. However, one limitation of including epigenetic features is that epigenetic data is not available for all cell types or species. This might cause decision challenges when trying to add these epigenetic features into learning model. To evaluate which combination of features is optimal for gRNA activity prediction, we generated a combined set of 2701 features including epigenetic markers (see method, feature set III). These features have been validated or hypothetically suggested to be important for CRISPR gRNA activity.

Next, we used minimal redundancy maximal relevance (mRMR) method to evaluate which features are important for gRNA activity prediction in our model. we choose a subset of the human Doench V1 data (NB4) as test data to do the feature selection (Table 1), because epigenetic features are available for the NB4 cell and can test the generalized performance of the selected features of whether IFS or not. In addition, performing feature selection and training in the same dataset can lead to overfitting. We applied the incremental feature selection (IFS) method to choose the optimum features for the dataset and model. First, we assumed that S_1_, S_2_ … S_N_ was in the set of S_i=_ [S_1_, S_2_ … S_N_]. Then, each feature set was used to train the model, and the feature set yielding the best performance was extracted. Based on the Bayesian Ridge Regression (BBR), we tested the optimal combination of features and dataset for gRNA activity prediction. Two best performing datasets were used: the normalized Hela dataset and the combined normalized Hela and HCT116 dataset. The model with the highest Spearman Correlation Coefficient (0.413) was achieved when 1839 features were used after IFS in feature set III of hct116 combine Hela dataset. Moreover, the Spearman Correlation Coefficient is 0.385 when 2029 features were used through IFS using the Hela dataset in feature set III. With the pre-feature set II (see method), we obtained 1163 and 777 features by IFS with the best Spearman correlation for the combined and Hela data, respectively (Figure 5a).

**Figure 5:**
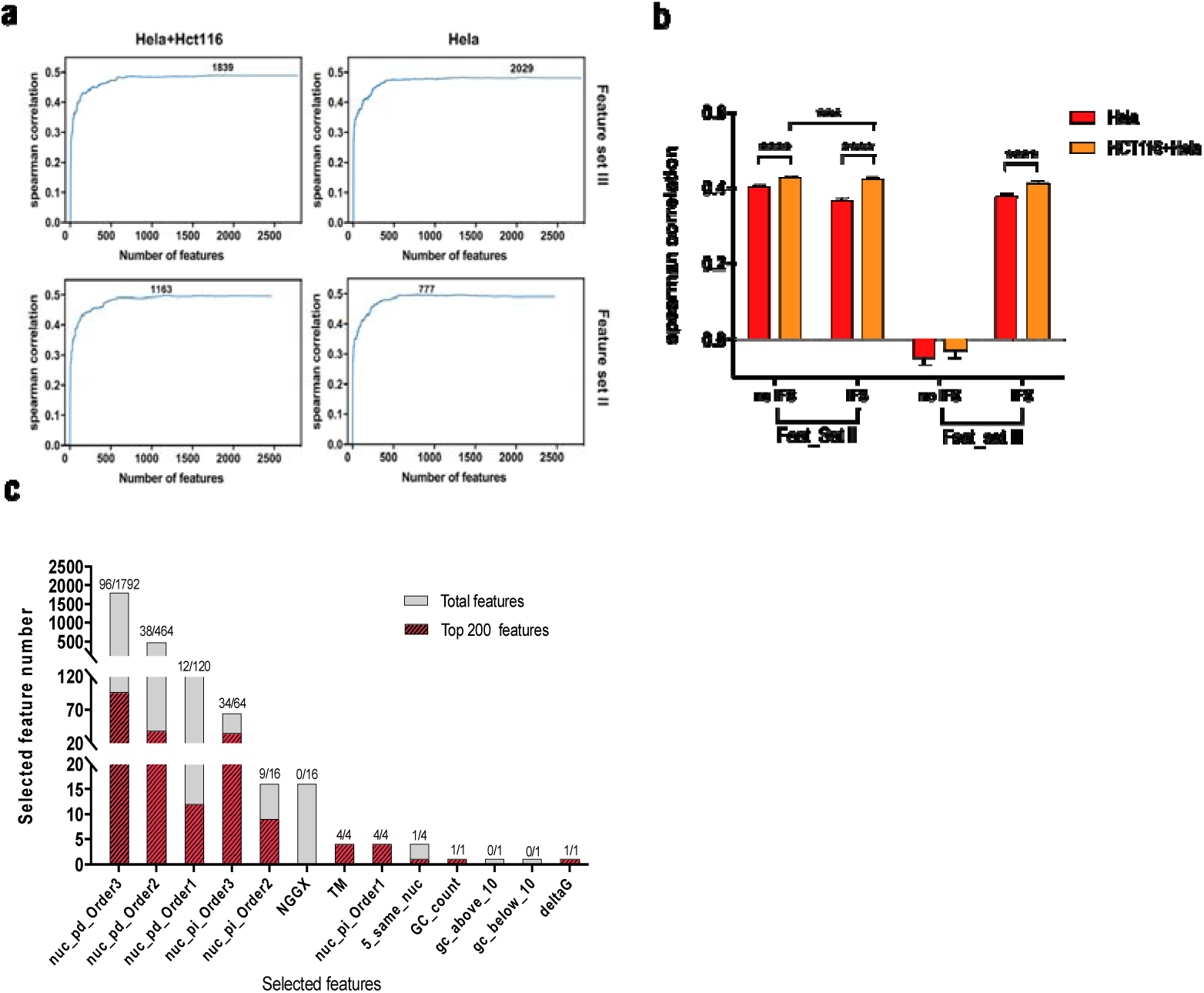
The optimized features selection result. a) The feature selection is conducted by IFS in feature set III and feature set II respectively for searching the best combination of feature in each optimized selected dataset. The upper two figures are the feature selection applied in feature set III, which results in 2,701 features in total, including pssm, disorder, epigenetic features; the two figures below are the feature selection used in the 2,488 features from feature set II. b) Evaluating the optimized models using the NB4 dataset. Two models are trained by feature set II is shown on the left side, while the ones trained by feature set III is shown on the right side. Each feature set is processed by incremental feature selection (IFS) compared with that of not feature selection. The number of features selected by IFS in feature set II are 777 and 1,163 in the model of Hela and Hct116+Hela, respectively. In feature set III, the features number are 2,029 and 1,839, respectively. c) The statistic of the top 200 optimized features selected by the NB4 cell type dataset after ranking by the algorithm of mRMR in the dataset of Hela using the whole 2,488 features in feature set II. Statistical significance is derived from t-test*. Asterisk* (*) indicates a *p* value less than 0.05, (**) less than 0.01, (***) less than 0.0001, and (****) less than 0.00001.

We also compare the IFS features to the non-IFS model by using 2488 of feature set II and 2701 of feature set III. Our results showed that the model trained by Hct116 + Hela performed the best in the NB4 test data without IFS used the feature set II (2488 features in total) (0.442) (Figure 5b). Also, we found that the hela+hct116 model outperforms the Hela model in the human dataset, so the feature selection is not necessary for the best performing model. For example, sgRNA Designer (Rule set II) picks up 13 types of features out of 627 features to train the GBRT regression model (Doench, et al., 2016) and sgRNA scorer utilizes 92 features without any selection to develop a classify model for stCas9 and spCas9 (Chari, et al., 2015).

Next, we evaluated which type of features (among these 2488 optimal feature sets trained with the combined hct116 and Hela datasets) plays the most decisive role in CRISPR gRNA activity. The GC count, nuc_pi_Order1, thermodynamic temperature, the secondary structure of the guide (deltaG) were regarded as the optimal features, because they all showed in the top 200 important features in the mrmr result (Figure 5c). The secondary structure of the guide (deltaG) affect CRISPR activity is consistent with our previous findings (Jensen, et al., 2017). Our featurization results suggest that the sequence composition, physicochemical characteristics greatly influence CRISPR gene editing activity. Our analysis also demonstrates that prediction software based on CNN such as DeepCRISPR (Chuai, et al., 2018) and CRISPRCpf1 (Kim, et al., 2018) can soundly predict the efficiency by extracting the feature using the convolution. When ranking was considered, the sequence thermodynamic temperature (TM) is ranked as the most important and decisive feature on gRNA activity. Previously studies have proven that unwinding was the primary determinant of CRISPR activity (Gong, et al., 2018) and Hui Peng et al. also observed the same outcome (Peng, et al., 2018). Among the top 20 most important features, the composition of continuous bases in specific position in the 20nt of sgRNA take the major importance in the sgRNA activity decision (80%) and the TM was rank as the most important feature, which mean position dependent and the TM the major role in the efficiency in the model trained by combined hct116+hela datasets.

### Develop new algorithms with improved general gRNA activity prediction

Current algorithms were trained with datasets from a specific cell type and frequently overfitting. This makes the algorithms incompatible with other cell type and species (Yan, et al., 2018). Labuhn et al. had previously proved that the state-of-art algorithm cannot be effectively generalized in small datasets (and) as the Spearman Correlation was just 0.2011 (Labuhn, et al., 2017). To develop an improved CRISPR gRNA activity prediction algorithm with better compatibility among different cell types and species (or generalization), we used the two best performing datasets (Hela and HCT116), the feature set II (see method without IFS), and BRR model to develop two generalization scores (GNL1.0 and GNL2.0). GNL1.0 was based on BBR trained with the normalized Hela dataset and feature set II. GNL2.0 was based on BBR trained with the combine Hela + HCT116 datasets and feature set II, whose datasets was seen as the best performance in the human dataset (Figure 5). The outcome of our study supports the result of Wilson et al. training on the dataset measured by sequencing results in more accurate activity predictions (Wilson, et al., 2018).

To evaluate the generalization, we compared GNL scores to seven algorithms (sgRNA Scorer, DeepCas9, sgRNA designer (rule set I), sgRNA designer (rule set II), SSC, CRISPR_Scan and Wu_CRISPR). Six datasets were used for evaluation (HEL (Labuhn, et al., 2017), Ciona (Gandhi, et al., 2016), *C. elegans* (Farboud and Meyer, 2015), *Drosophila* (Ren, et al., 2014), Z_fish_VZ (Varshney, et al., 2015), Z_fish_GZ (Gagnon, et al., 2014)). The GNL Scores showed the best generalized prediction outcome (Figure 6). The CRISPR-scan algorithm performed well in the zebrafish dataset, but gave a poor prediction outcome in human CRISPR gRNA activity. Previously, using the Hela dataset, it was found that the gRNA Designer (rule set II) performs better than the gRNA designer (rule set I), gRNA Scorer, SSC, and CRISPR-Scan (Chuai, et al., 2018). Consistent with that, our analysis showed that the gRNA Designer (rule set II) performs better than most of the algorithms evaluated in the human HEL dataset. Most importantly, considered all six testing datasets, the GNL1.0 and GNL2.0 have the best generalized prediction. The GNL2.0 is ranked as the best algorithms in predicting the human HEL dataset. Taken together, we have developed two CRISPR gRNA activity prediction algorithms with improved generalization and human gRNA activity prediction. The GNL scores can be freely accessed in GitHub (https://github.com/TerminatorJ/GNL_Scorer).

**Figure 6:**
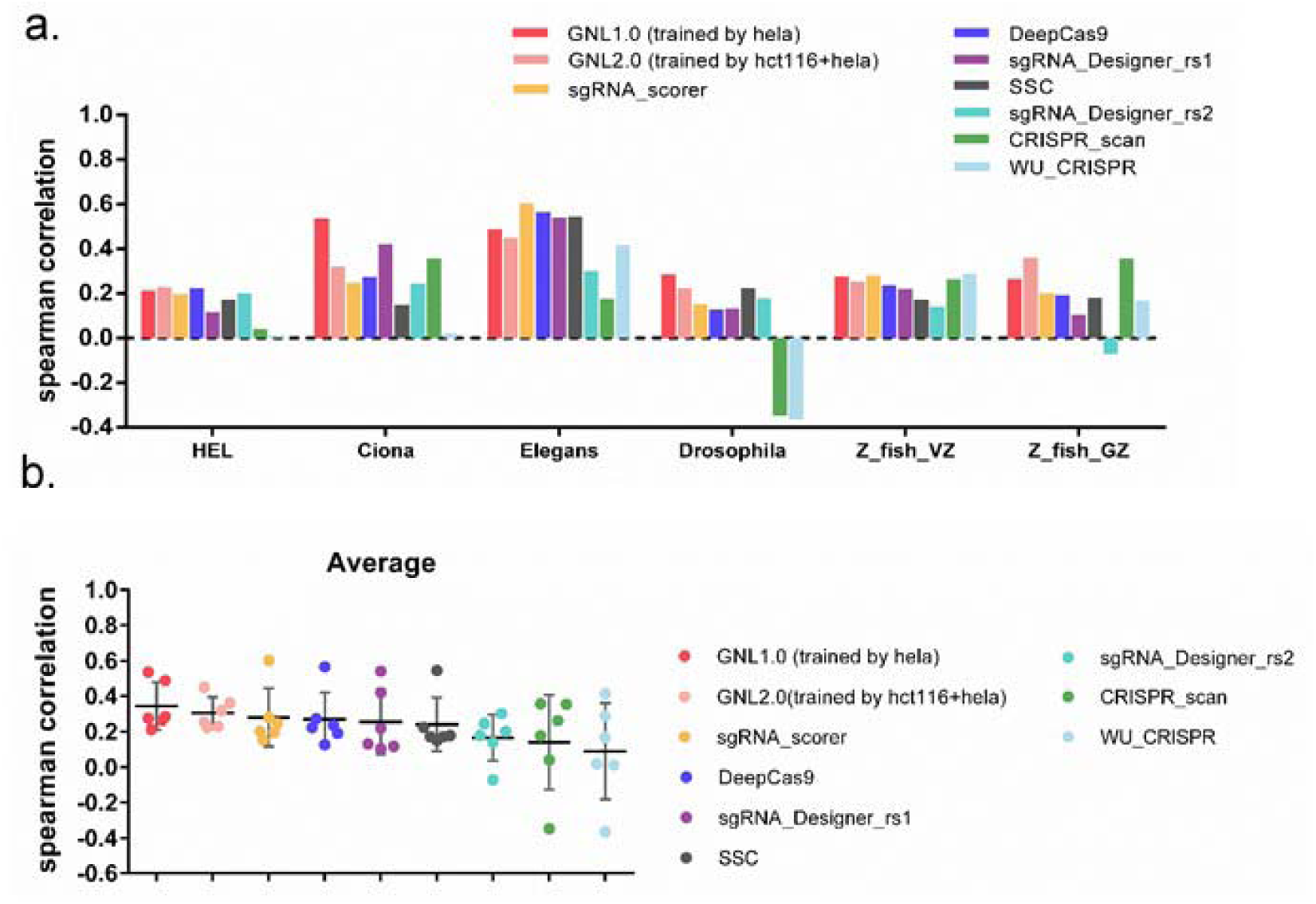
Comparison of model performance. GNL1.0 (general scorer 1.0) and GNL2.0 (general scorer 1.0) are models we training in this paper, using the best performance datasets (hela or hela+hct116) and the optimized feature set. These two models outperform the seven other algorithms in different species datasets include human.

## DISCUSSION

In this study, using combined normalized datasets, we have developed an improved model for predicting (or an improved prediction model) for CRISPR gRNA activity. One advantage of our model is that it provides a better generalized prediction compared to the existing algorithms. This provides a solution to the biggest disadvantage that most of the machine-learning based software is encountering. However, as shown in our analysis of the 13 datasets, there exists a great batch effect and variability of the CRISPR gRNA activity. This might be caused by many factors, such as the experimental condition, the method of gRNA activity measurement, the endogenous DNA repair machinery. The most basic principle of CRISPR gene editing is the precision introduction of a DNA double-strand break (DSB) to the target site, and repair of the DSB by the endogenous DNA repair machinery (Jiang and Doudna, 2017). In mammalian cells, the DSBs are predominantly repaired by the error-prone non-homologous end joining (NHEJ) and the microhomology-mediated end joining (MMEJ) pathways (Yao, et al., 2017), which will cause small deletions or insertions to the target site (also known as indels).

However, due to the impracticability of measuring the indel frequency caused by thousands of individual gRNA, all current approaches used to quantify CRISPR gRNA activity are based on functional readouts such as quantification of fluorescent inactivation by FACS (Doench, et al., 2014), number of resistant clones (Doench, et al., 2016), or quantification based on expression (Hart, et al., 2015). This partially explains the poor cross-dataset validation and correction of the different gRNA activity prediction algorithms, as well as the poor generalized cross-species prediction. To partially overcome this limitation, we in this study applied data normalization, dataset filtering, and dataset combination to develop an algorithm with improved generalized CRISPR gRNA activity prediction. Using the most representative detect datasets, we developed our own model based on the Bayesian Ridge Regression by machine learning. However, the current datasets are still not sufficient enough to build a strong prediction model with high correlations. The batch-effect contributes greatly to the generalization of different models. Thus, there is still a great need to build a benchmark and large-scale approach to experimentally measure the CRISPR activity from a large number of gRNAs.

Both the Spearman Correlation and ROC-AUC curve are typical measurements for evaluating the performance of CRISPR gRNA gene editing activity prediction models. We only used the Spearman Correlation Coefficient in this study, because the ROC-AUC curve was previously shown to be less informative for those RES and FC datasets generated by flowcytometry and resistant assay for gene KO (Fusi, et al., 2015).

Another improvement of our model is featurization. Factors, also known as features, systematically and coherently decides why CRISPR-Cas9 can achieve high targeting efficiency at one locus but not the other. The majority of these decisive features are sequence dependent, while there are still many other features such as chromatin accessibility, DNA repair machinery, epigenetic markers, expression states that are not related to sequences. We had previously experimentally identified some of these important features (Jensen, et al., 2017). However, there might be still many unknown features, sequence-dependent and sequence-independent features that need to be identified in the future. Nevertheless, using machine learning in this study we identified and proved that physicochemical characteristics, sequence composition, the epigenetic features and gRNA secondary structure greatly affect CRISPR gRNA gene editing efficiency. Taken together, we have developed an improved CRISPR gRNA activity prediction algorithms which will contribute to the current in silico design of CRISPR gRNA and will promote the appropriate applications of the CRISPR gene editing technology.

## ACKNOWLEDGEMENTS

We thank the Dr. Shan and Dr. Bai for many instructive advices in the statistics and algorithms respectively. And Haeussler who shared the data they collected which save the time of us for initially data processing.

## FUNDING

This work was supported by the Lundbeck Foundation (R219–2016-1375, R173–2014-1105), the Danish Research Council for Independent Research (DFF–1337–00128), the Sapere Aude Young Research Talent Prize (DFF-1335–00763A), the Innovation Fund Denmark (BrainStem), and Aarhus University Strategic Grant (AU-iCRISPR). Y.L. is also supported by Guangdong Provincial Key Laboratory of Genome Read and Write (No. 2017B030301011).

