## supplementary Data for "CRISPR-GNL: an improved model for predicting CRISPR activity by machine learning and featurization"

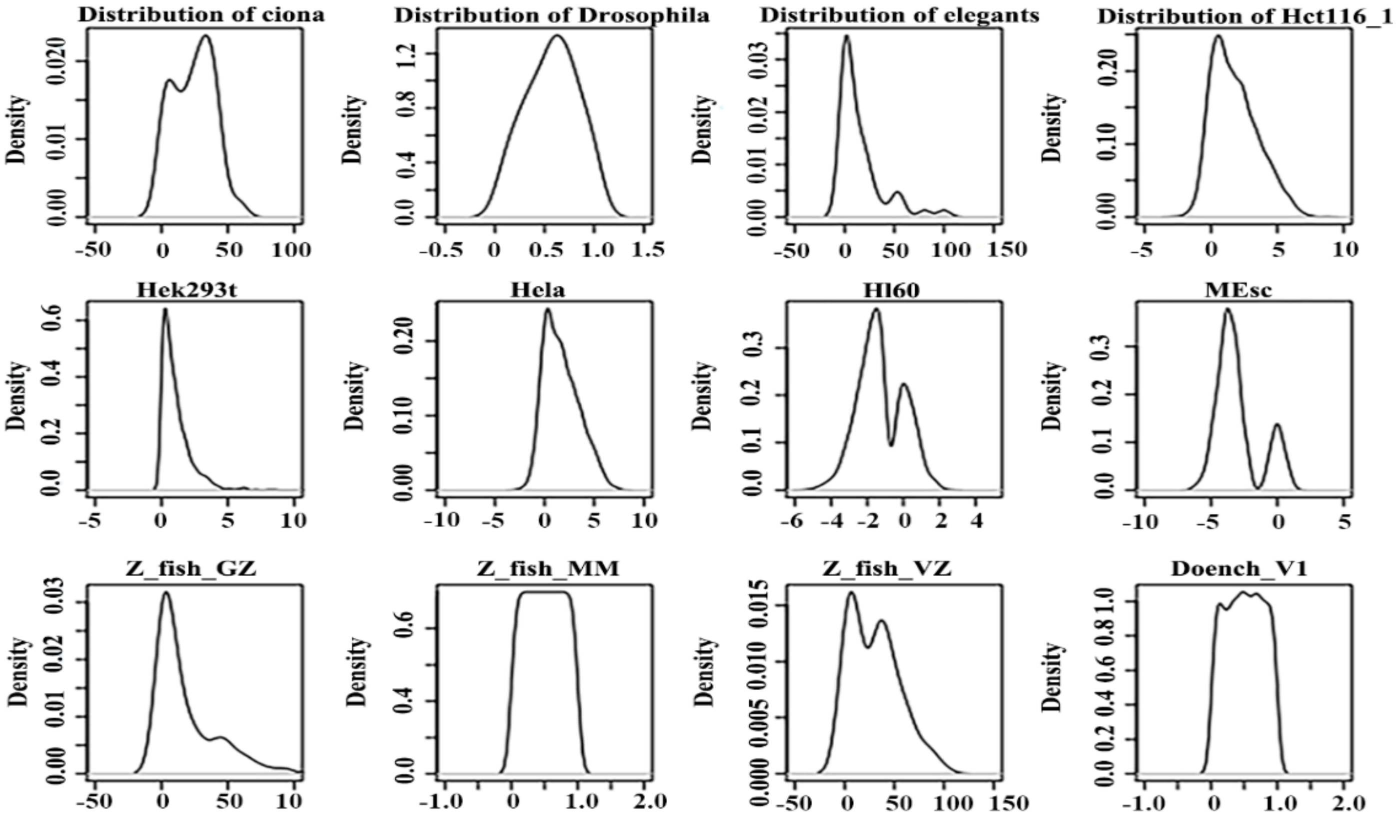


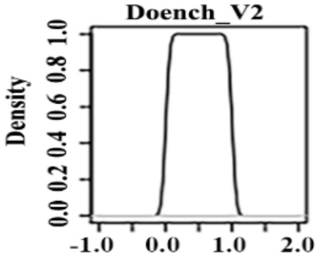


**Figure S1: 13 raw datasets before normalization.** The density plots show the distribution of 13 initial datasets. Hct116 means the hct116 cell type data library 1 from Hart et al. Z_fish_GZ; Z_fish_MM; Z_fish_VZ are three zebrafish datasets from Gagnon et al.[24], Moreno-Mateos et al.[16], Varshney et al.[25].

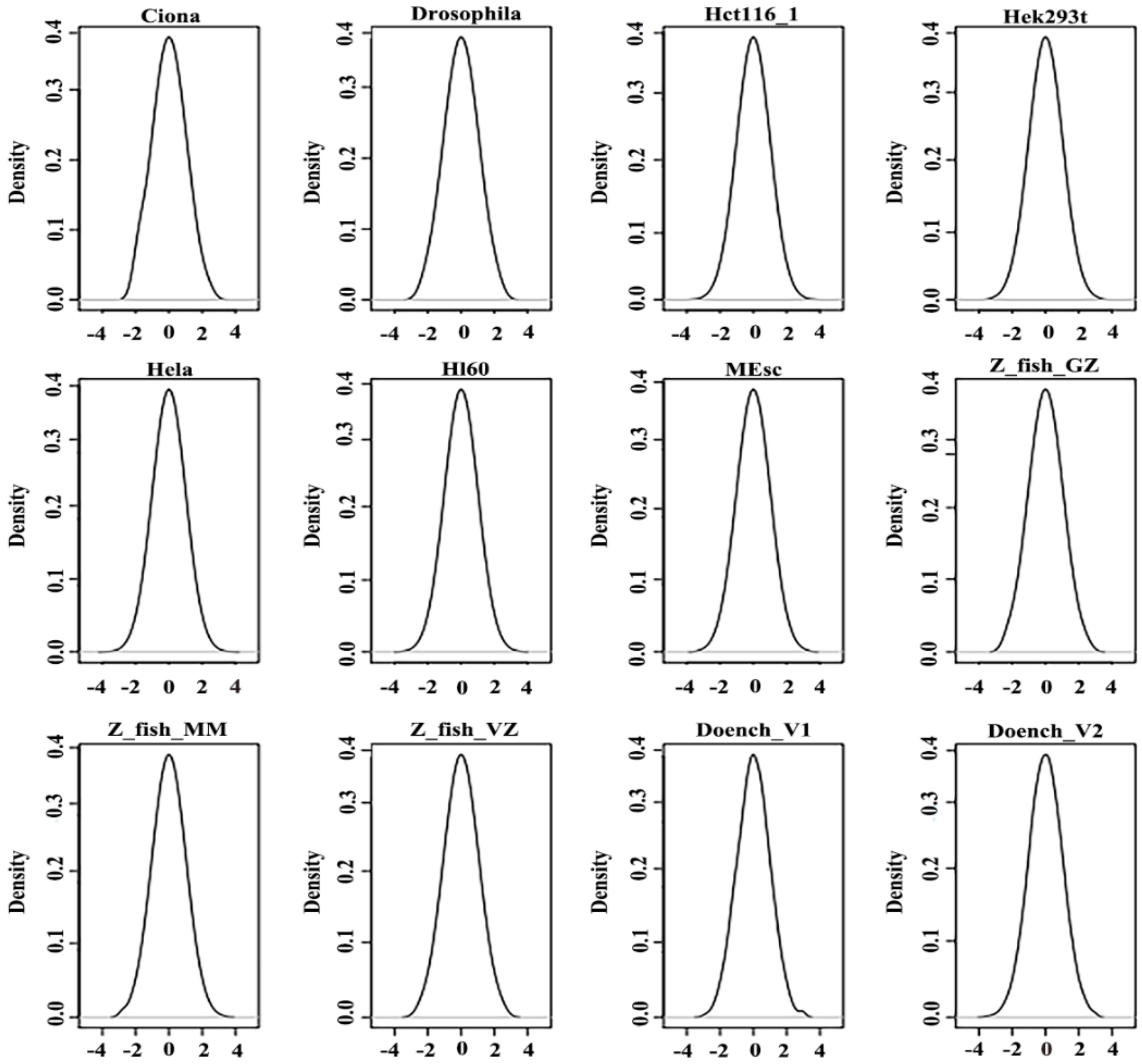


Figure S2: 12 datasets after normalization. We normalize these datasets to the range of -4 to 4.

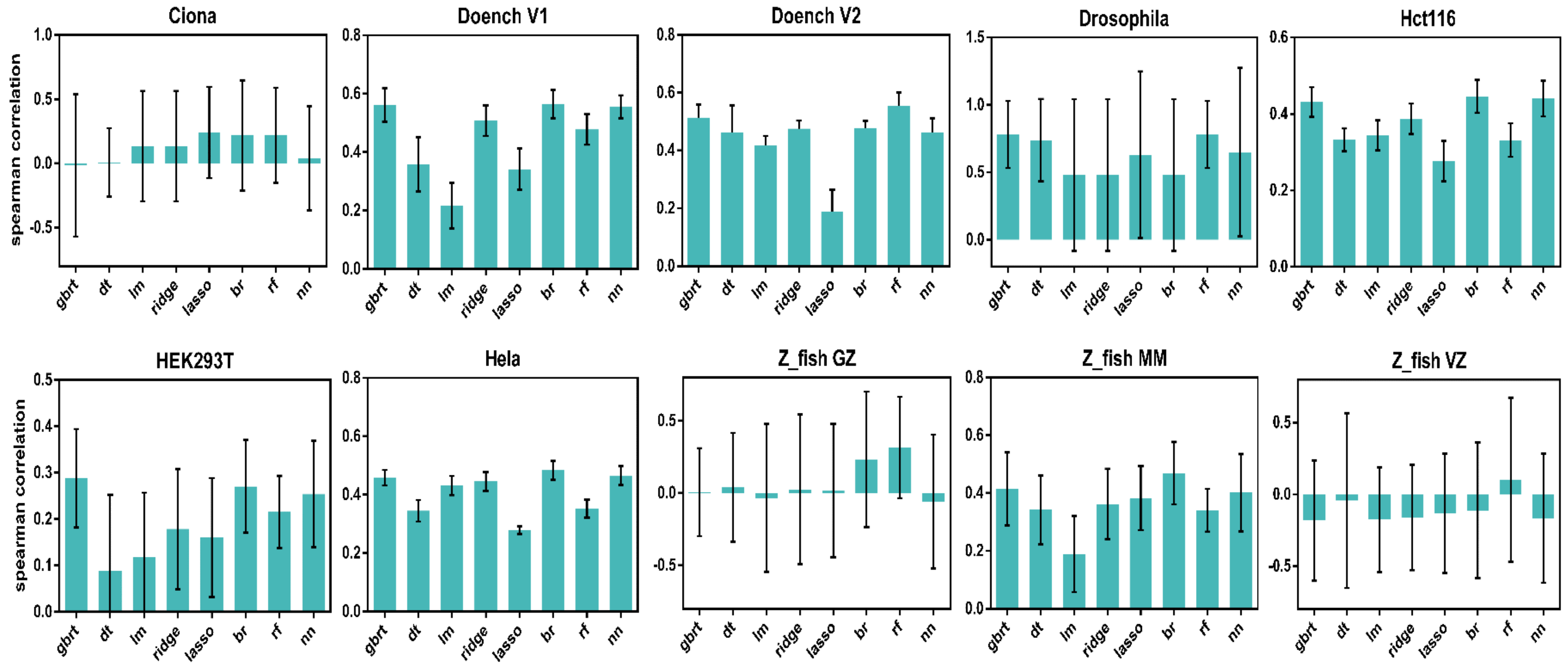


**Figure S3: 10 normalized datasets are trained on eight different regression models.** Each bar shows the mean value of Spearman correlation trained by each dataset. Error bars show the s.d. across sgRNA with a 10-fold cross validation approach. For each dataset, we regard the highest Spearman correlation and minimum standard deviation model as the best model.

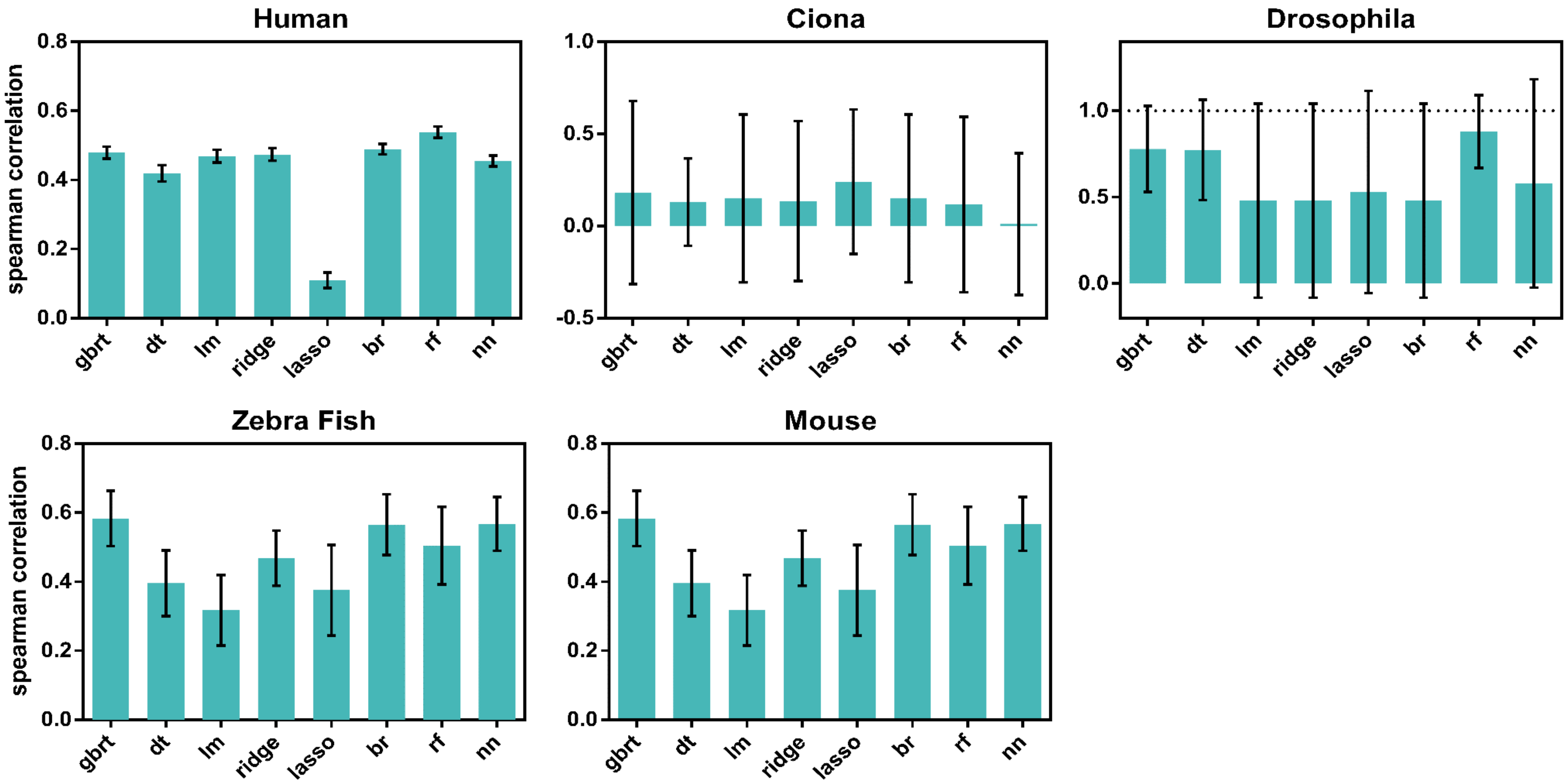


**Figure S4: 5 normalized species datasets are trained on eight different regression models.** Each bar shows the mean value of Spearman correlation. Error bars show the s.d. across sgRNA with a 10-fold cross validation approach. For each dataset, we regard the highest Spearman correlation and minimum standard deviation as the best model.
